# Rapid high-resolution structure analysis of small, biotechnologically relevant enzymes by cryo-electron microscopy

**DOI:** 10.1101/2021.06.15.448552

**Authors:** Nicole Dimos, Carl P.O. Helmer, Andrea M. Chánique, Markus C. Wahl, Robert Kourist, Tarek Hilal, Bernhard Loll

## Abstract

Enzyme catalysis has emerged as a key technology for developing efficient, sustainable processes in the chemical, biotechnological and pharmaceutical industries. Plants provide large and diverse pools of biosynthetic enzymes that facilitate complex reactions, such as the formation of intricate terpene carbon skeletons, with exquisite specificity. High-resolution structural analysis of these enzymes is crucial to understand their mechanisms and modulate their properties by targeted engineering. Although cryo-electron microscopy (cryo-EM) has revolutionized structural biology, its applicability to high-resolution structure analysis of comparatively small enzymes is so far largely unexplored. Here, we show that cryo-EM can reveal the structures of ~120 kDa plant borneol dehydrogenases at or below 2 Å resolution, paving the way for the fast development of new biocatalysts that provide access to bioactive terpenes and terpenoids.

## Introduction

Despite the stunning recent success of single-particle cryo-EM in the structure analysis of many large molecular machines, comparatively small proteins remain a major challenge for this technique^1–3^. To date, the highest resolution achieved by cryo-EM is 1.14 Å for the highly symmetric 480 kDa protein, apoferritin^4^. Presently, the cryo-EM structures of only four macromolecular complexes smaller than 120 kDa have been reported at a resolution better than 3.0 Å (**Supplementary Table 1**). Due to limited structural data and frequently insufficient understanding of the molecular basis of enzyme catalysis, protein engineering is still mainly relying on combinatorial approaches for enzyme engineering such as random or saturation mutagenesis. An expansion of the scope of single particle analysis towards the rapid elucidation of the structures of smaller proteins has a tremendous potential to increase the rational element of protein engineering.

Enzyme catalysis offers an efficient and sustainable alternative to traditional chemical synthesis, as biocatalysts harbor excellent selectivity and work under mild reaction conditions. Today, enzymes are widely used in the chemical and pharmaceutical industries, and the application of biocatalysts for the manufacture of chemicals from renewable resources is a rapidly growing field. However, due to limited structural data and, as a consequence, insufficient understanding of the molecular basis of enzyme catalysis, rational improvement of biotechnologically relevant enzymes is severely hampered. Protein engineering relies mainly on directed evolution or semi-rational approaches. While often successful, these methods require the implementation of high-throughput screenings and a substantial effort in terms of laboratory work, leading to long time-to-market horizons. This situation could be alleviated by expanding the scope of cryo-EM towards the rapid elucidation of high-resolution structures of smaller proteins.

A particularly interesting application of enzyme catalysis is the synthesis and modification of bioactive terpenes and terpenoids. With more than 50,000 different structures, terpenes are a structurally and functionally extraordinarily diverse group of natural products^5^. The outstanding selectivity of the enzymes involved in the formation of terpene carbon skeletons^6^, their primary functionalization^7^ and further derivatization^8^, could enable the formation of a myriad of new terpene derivatives with diverse, interesting properties for the food and pharmaceutical industries *via* environmentally friendly catalytic processes^5, 9^. To this end, a detailed understanding of the molecular basis of the enzymes’ reaction mechanisms and selectivity is required.

Bornane-type monoterpenoids, such as borneol, isoborneol and camphor, are found in essential oils from plants and are used in traditional medicine and cosmetics^10^. Racemic borneol, isoborneol and camphor are currently produced from α-pinene, a side product of cellulose production. Essential oils from plants are often enriched in one of the enantiomers of these compounds, indicating the potential presence of highly stereoselective borneol dehydrogenases (BDH). An enzymatic route towards pure enantiomers using enantioselective dehydrogenases would be highly desirable to avoid the labor-intensive and expensive extraction of pure enantiomers from plants.

BDHs belong to the family of short-chain dehydrogenases-reductases (SDR)^11^. Members of this enzyme class have a TGxxx[AG]xG NAD^+^-binding motif and a YxxxK active site motif^12, 13^ and form dimers or tetramers. Some BDHs have a high, two-fold stereoselectivity in the conversion of chiral monoterpenoids^11, 14, 15^ by preferring one of the two substrates enantiomers and forming a stereo-center by the asymmetric reduction of the diastereotopic keto-group. The selective oxidation of (+)-borneol to (+)-camphor by a partially purified BDH from *Salvia officinalis L.* was first described by Croteau and co-workers^14^. Later, the isolation and purification of two BDHs by us confirmed the high stereoselectivity of the enzymes (**Fig. 1a,b, and c**)^11, 15^.

**Fig. 1.**
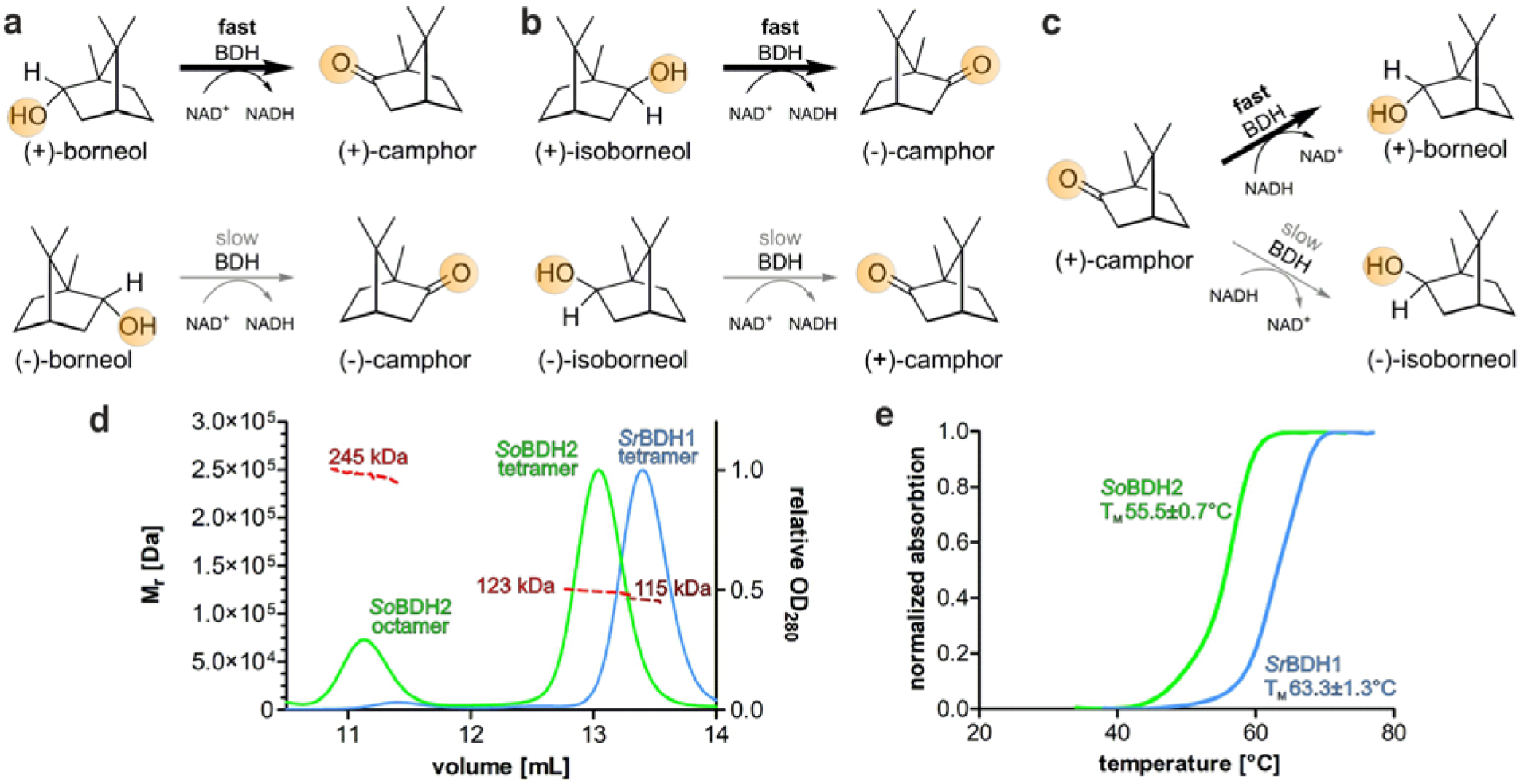
Reaction schemes and enzyme characterization of *So*BDH2 and *Sr*BDH1. The here discussed BDHs preferentially convert the (+)-enantiomers of borneol and isoborneol. *Sr*BDH1 is highly selective for both alcohols and catalyzes the reduction of (+)-camphor. *So*BDH2 is highly selective for borneol. *Ps*BDH shows merely a slight selectivity for both borneol and isoborneol and is capable of catalyzing the reduction of camphor^16^. **a**,**b** reaction schemes of *Sr*BDH1 and *So*BDH2 in the enantiospecific oxidation of *rac-*borneol (**a**) and *rac-*isoborneol (**b**). **c** reduction of (+)-camphor. **d** SEC/MALS analysis of *Sr*BDH1 (blue) and *So*BDH2 (green). For *Sr*BDH1 a single peak is observed consistent with a tetramer (theoretical Mr 120 kDa). The first peak in the chromatogram of *So*BDH2 corresponds to an octamer (theoretical Mr 258 kDa) and a second peak corresponding to tetramer (theoretical Mr 129 kDa). Brown (*Sr*BDH1) and red (*So*BDH2) curves, refractive index signals. **e** Differential scanning fluorometry reveals a significantly lower T_M_ for *So*BDH2 (55.5±0.7°C) compared to *Sr*BDH1 (63.3±1.3°C).

The understanding of terpene formation on a structural and mechanistic basis is important for the engineering of biosynthetic pathways for the formation of new terpenoids^17^. Unfortunately, the difficulty to produce enzymes from higher organisms in bacteria, and often their limited stability, make structure determination by classical crystallization very challenging. In order to obtain a structure under these circumstances, approaches such as truncation and homology modeling are utilized, with both having a limited informational value for mechanistic studies and enzyme engineering. In the particular case of BDHs, only two crystal structures have been reported to date, *i.e.* of the unselective bacterial BDH from *Pseudomonas* sp. TCU-HL1 (*Ps*BDH; PDB ID 6M5N^16^) and the enantioselective BDH from *S. rosmarinus* (*Sr*BDH1)^11^. Although the structure analysis of *Sr*BDH1 allowed us to identify a hydrophobic pocket that discriminates the monoterpenol isoborneol, structures of additional BDHs, e.g. from *S. officinalis* (*So*BDH2), are required to rationalize the selectivity of the enzymes towards (+)-borneol. Here, we report the rapid determination of the structures of two stereoselective dehydrogenases, *Sr*BDH1 and *So*BDH2, by single-particle cryo-EM.

## Results

### High resolution cryo-EM structure of *So*BDH2

*Sr*BDH1 and *So*BDH2 exhibit 44 % and 60 % sequence identity and similarity, respectively. We produced *So*BDH2 with a N-terminal His6-tag (32.2 kDa theoretical molecular weight (Mr)) in *E. coli* and prepared the protein at high purity. Size exclusion chromatography coupled to multi-angle light scattering (SEC/MALS) revealed two distinct species (**Fig. 1d**), corresponding to an octameric and a tetrameric assembly.

Encouraged by the possible occurrence as octameric assembly, we considered cryo-EM as powerful method to dissect structural heterogeneity, and prepared cryo-EM grids. Imaging was conducted on a Titan Krios 300 kV TEM equipped with a Falcon 3EC detector operated in counting mode. We aligned the instrument thoroughly and aimed to maximize beam coherence by choosing a 50 μm *C*2 aperture. To optimize the *C*2 intensity and stigmation, we used the ronchigram method on a Volta phase plate (VPP^18^). The VPP was only used for alignment and retracted during data acquisition. A total of 1,439 micrographs were acquired and subjected to motion correction and CTF estimation. From 1,551,724 particle images initially picked, 173,781 particle images were selected by iterative 2D and 3D classification cycles for homogeneous 3D refinement (**Supplementary Fig. 1** and **Supplementary Fig. 2**). Although we had observed a fraction of octamers in solution (**Fig. 2c**), 3D refinement yielded merely a tetrameric structure (**Supplementary Fig. 2**); we also failed to detect octamers in negative stain EM.

**Fig. 2.**
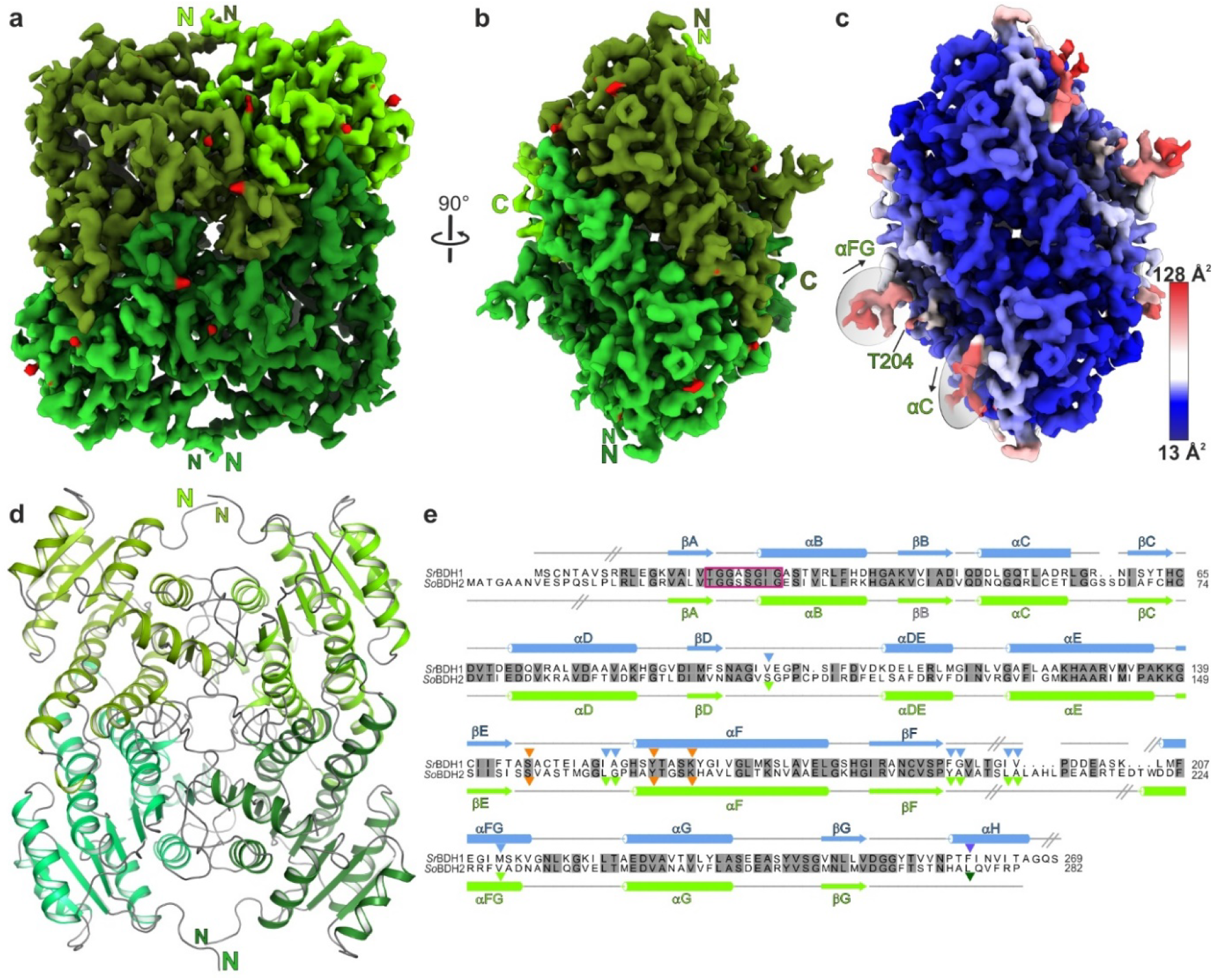
cryo-EM structure of *So*BDH2. **a** Tetrameric assembly of *So*BDH2. Density is shown for each of the four protomers in different green shades. In red, locations of very well defined water molecules. **b** Same color-coding as in a, but rotated by 90 °. **c** grouped B-factor mapped on the density. Color-gradient from blue to red, corresponding to increasing B-factors. Regions with high B-factors are highlighted with grey ellipses and labelled according to the assigned secondary structure. **d** *So*BDH2 structure in cartoon representation. Same view and color-coding as in panel a. **e** Structure-based sequence alignment of *Sr*BDH1 (GeneBank ID MT857224) and *So*BDH2 (GeneBank ID MT525099) as obtained by cryo-EM. Secondary structure elements are drawn above the alignment for *Sr*BDH1 and below for *So*BDH2 with α-helices depicted as cylinders and β-strands as arrows. Grey, inclined lines indicated sections of the structures, since they were not resolved in the reconstruction and could not be modelled. Orange triangles indicate the catalytic motif. Amino acids lining the putative active site of *Sr*BDH1, based on its crystal structure (PDB ID 6ZYZ) with bound NAD^+^, are indicated by blue triangles and as dark blue triangle if derived from another *Sr*BDH1 monomer within the tetramer. The dark green triangle marks a residue derived from another protomer of *So*BDH2 that completes the active site. Grey shaded amino acids are identical. The TGxxx[AG]xG NAD^+^-binding motif, between βA and αB is indicated with a magenta rectangle.

After application of global and local CTF refinement, particle-based local motion correction and NU refinement within the cryoSPARC framework^19^ a final gold-standard resolution of 2.04 Å was obtained. This, to the best of our knowledge, is the highest reported resolution of a sub 200 kDa protein solved by single-particle cryo-EM. The obtained cryo-EM density reflects the nominal resolution, as individual side chains could be unambiguously identified and built. Moreover, spherical density clearly indicated the presence of water molecules. Given the high resolution of our cryo-EM map (**Fig. 2a,b** and **Table 1**), we tested how automated model building programs ARP/wARP^20^, phenix.map_to_model^21^ and Buccaneer^22^ would perform. The programs were run with the recommended standard settings and the results are summarized in **Supplementary Table 2**. All programs managed to fit large portions of the protein sequence to the density (82 – 92 %) with ARP/wARP outperforming the other two programs. We manually completed the initial ARP/wARP model. Spherical density regions clearly indicated water molecules and well-defined water molecules were automatically placed with COOT^23^. The quality of the density allowed modelling of 50 double conformations of amino acid side chains and the localization of 268 water molecules. The final model exhibits an excellent fit to the density with mask/volume correlation coefficients of 0.86/0.83 (**Table 1**).

**Table 1.**
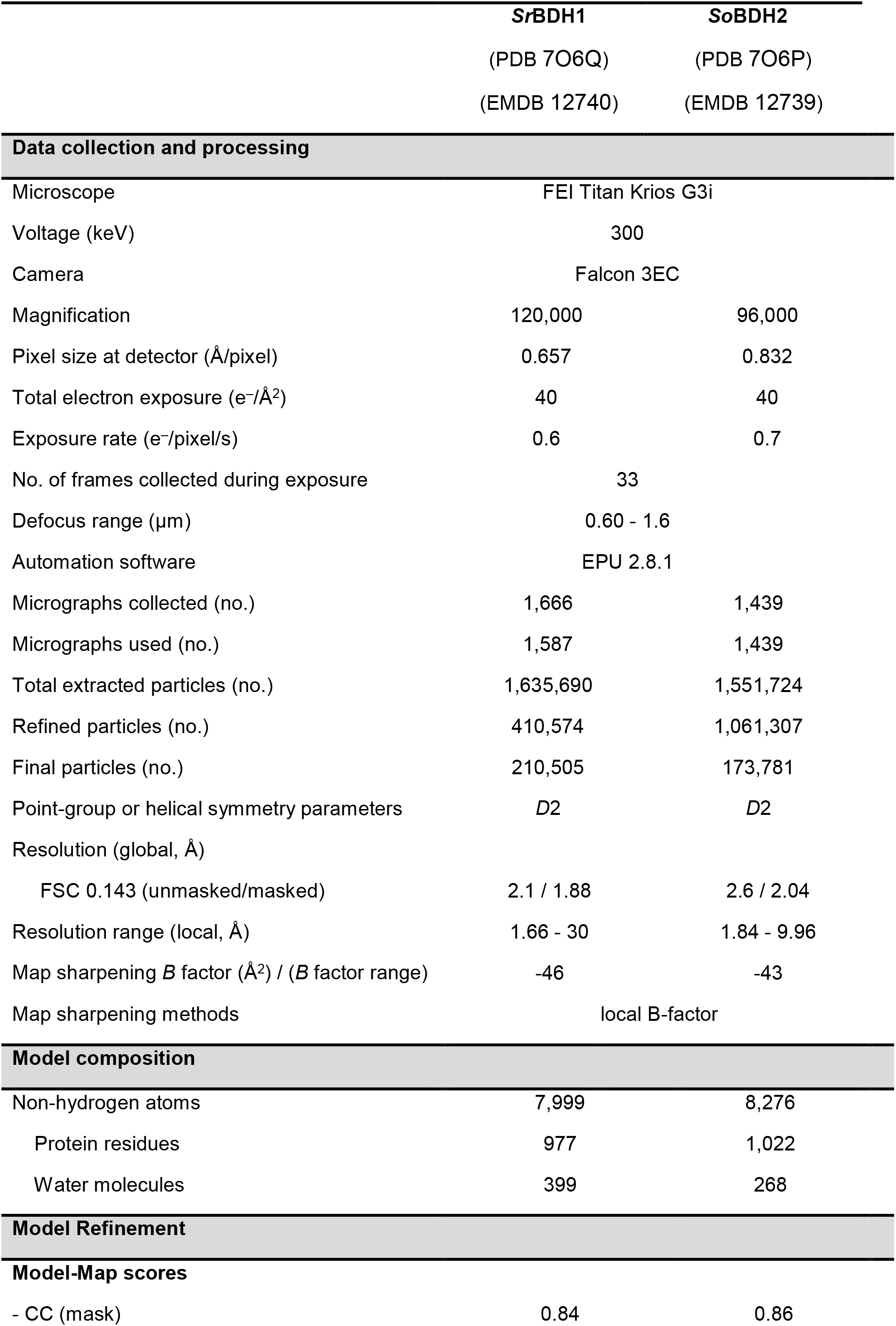
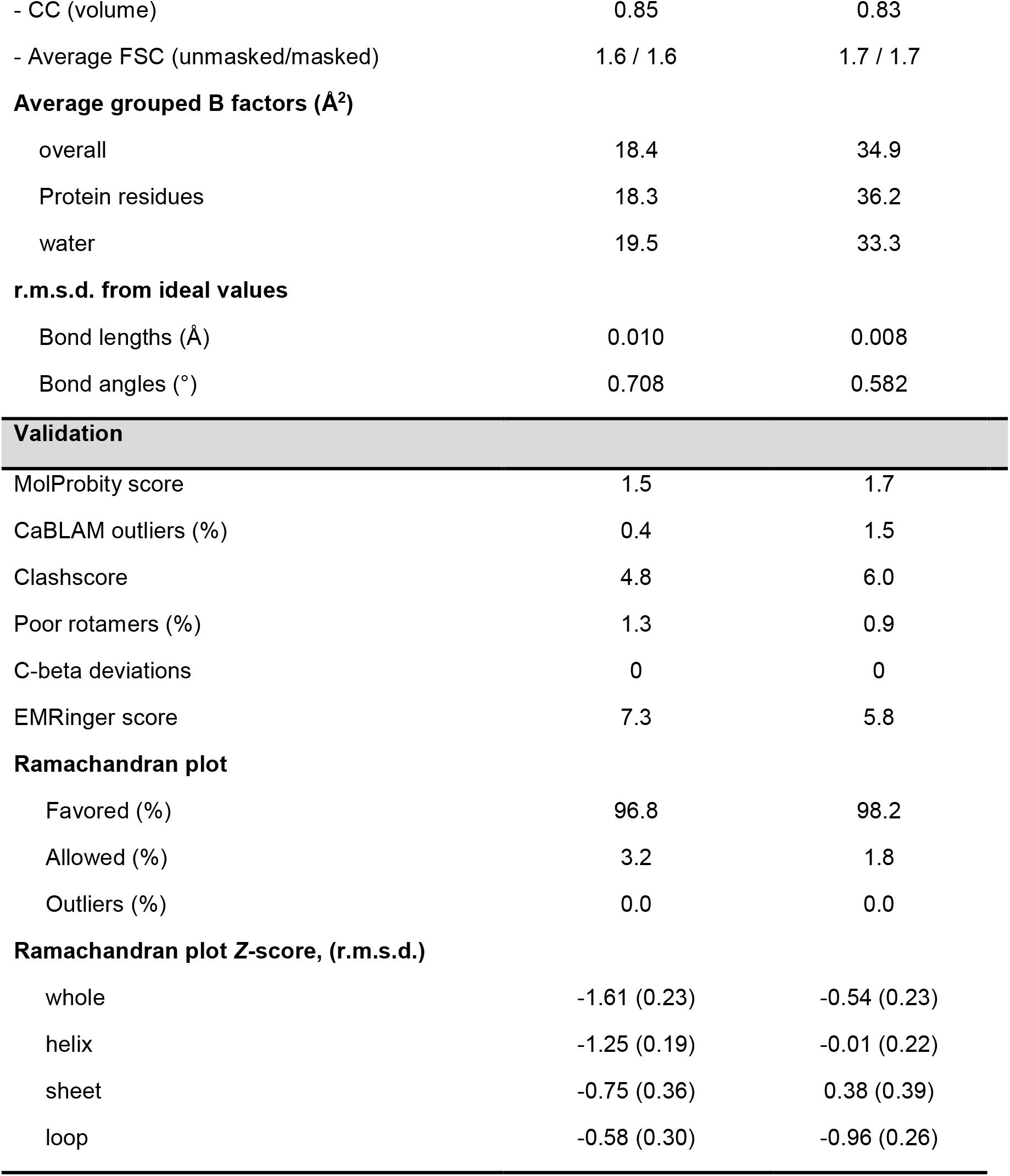
cryo-EM data collection, refinement, and validation statistics

|  | SrBDH1 | SoBDH2 |
| --- | --- | --- |
|  | (PDB 7O6Q) | (PDB 7O6P) |
|  | (EMDB 12740) | (EMDB 12739) |
| Data collection and processing |  |  |
| Microscope | FEI Titan Krios G3i |  |
| Voltage (keV) | 300 |  |
| Camera | Falcon 3EC |  |
| Magnification | 120,000 | 96,000 |
| Pixel size at detector (Å/pixel) | 0.657 | 0.832 |
| Total electron exposure (e<sup>-</sup>/Å<sup>2</sup>) | 40 | 40 |
| Exposure rate (e<sup>-</sup>/pixel/s) | 0.6 | 0.7 |
| No. of frames collected during exposure | 33 |  |
| Defocus range (µm) | 0.60 - 1.6 |  |
| Automation software | EPU 2.8.1 |  |
| Micrographs collected (no.) | 1,666 | 1,439 |
| Micrographs used (no.) | 1,587 | 1,439 |
| Total extracted particles (no.) | 1,635,690 | 1,551,724 |
| Refined particles (no.) | 410,574 | 1,061,307 |
| Final particles (no.) | 210,505 | 173,781 |
| Point-group or helical symmetry parameters | *D*2 | *D*2 |
| Resolution (global, Å) |  |  |
| FSC 0.143 (unmasked/masked) | 2.1 / 1.88 | 2.6 / 2.04 |
| Resolution range (local, Å) | 1.66 - 30 | 1.84 - 9.96 |
| Map sharpening *B* factor (Å<sup>2</sup>) / (*B* factor range) | -46 | -43 |
| Map sharpening methods | local B-factor |  |
| Model composition |  |  |
| Non-hydrogen atoms | 7,999 | 8,276 |
| Protein residues | 977 | 1,022 |
| Water molecules | 399 | 268 |
| Model Refinement |  |  |
| Model-Map scores |  |  |
| - CC (mask) | 0.84 | 0.86 |
| - CC (volume) | 0.85 | 0.83 |
| - Average FSC (unmasked/masked) | 1.6 / 1.6 | 1.7 / 1.7 |
| **Average grouped B factors (Å<sup>2</sup>)** |  |  |
| overall | 18.4 | 34.9 |
| Protein residues | 18.3 | 36.2 |
| water | 19.5 | 33.3 |
| **r.m.s.d. from ideal values** |  |  |
| Bond lengths (Å) | 0.010 | 0.008 |
| Bond angles (°) | 0.708 | 0.582 |
| **Validation** |  |  |
| MolProbity score | 1.5 | 1.7 |
| CaBLAM outliers (%) | 0.4 | 1.5 |
| Clashscore | 4.8 | 6.0 |
| Poor rotamers (%) | 1.3 | 0.9 |
| C-beta deviations | 0 | 0 |
| EMRinger score | 7.3 | 5.8 |
| **Ramachandran plot** |  |  |
| Favored (%) | 96.8 | 98.2 |
| Allowed (%) | 3.2 | 1.8 |
| Outliers (%) | 0.0 | 0.0 |
| **Ramachandran plot Z-score, (r.m.s.d.)** |  |  |
| whole | -1.61 (0.23) | -0.54 (0.23) |
| helix | -1.25 (0.19) | -0.01 (0.22) |
| sheet | -0.75 (0.36) | 0.38 (0.39) |
| loop | -0.58 (0.30) | -0.96 (0.26) |

As already observed for the crystal structures of *Sr*BDH1 and *Ps*BDH, the *So*BDH2 homo-tetramer exhibits *D*2 symmetry (**Fig. 2** and **Supplementary Fig. 1**). The protomers adopt a Rossmann-like fold^24^, required for binding of the NAD^+^ cofactor. Twelve N-terminal residues and the preceding His6-tag lack density (**Fig. 2e**). The high resolution allowed us to locate 268 water molecules and 50 alternative side chain conformations. Very weak and fragmented density is observed for *So*BDH2 residues Q52 to G65 that fold into α-helix αC (**Fig. 2e**), reflected in elevated B-factors (**Fig. 2c**). Moreover, the region from S205 to E218 is not resolved in the density and has not been modelled (**Fig. 2e**) in agreement with the observation that we could not observe any density for the NAD^+^ cofactors in their binding pockets. The latter observation is in agreement with the apo state crystal structure of *Sr*BDH1. However, the crystal structure of apo*-Sr*BDH1 could be only obtained after co-crystallization with the substrate (+)-borneol, led to reduction of NAD^+^ and release of product and cofactor. Loss of the cofactor could not be prevented by adding a threefold molar excess of NAD^+^ to *So*BDH2 before size exclusion chromatography. While the loss of NAD^+^ may have occurred during vitrification of the cryo-EM sample, in the NAD^+^-bound crystal structures of *Sr*BDH1 the cofactor binding site is stabilized by crystal contacts, suggesting that under the crystallization conditions the NAD^+^ binding site is artificially stabilized to prevent release of the co-factor.

### Active site of *So*BDH2

Despite the absence of NAD^+^, the spatial arrangement of the catalytic S156, K169, Y173 motif (**Fig. 2e**) is maintained in *So*BDH2 compared to *Sr*BDH1•NAD^+^ ^11^ (**Fig. 3**). The lysine residue in concert with the positively charged nicotinamide lower the pK_a_ value of the tyrosine, which acts as catalytic acid/base. The serine residue is involved in stabilization and polarization of the carbonyl function of the substrate^25^. As in *Sr*BDH1, the substrate-binding niche is very hydrophobic, but decorated by different amino acid residues. Moreover, both enzymes have in common that the C-terminus of another protomer completes their active site pocket (**Fig. 3**). Notably, the C-terminus of *So*BDH2 adopts a coiled-coil structure in contrast to the C-terminal α-helix αH in *Sr*BDH1 (**Supplementary Fig. 5a**), but both *Sr*BDH1 F260 and *So*BDH2 L277 reside in the same position (**Fig. 3b**).

**Fig. 3.**
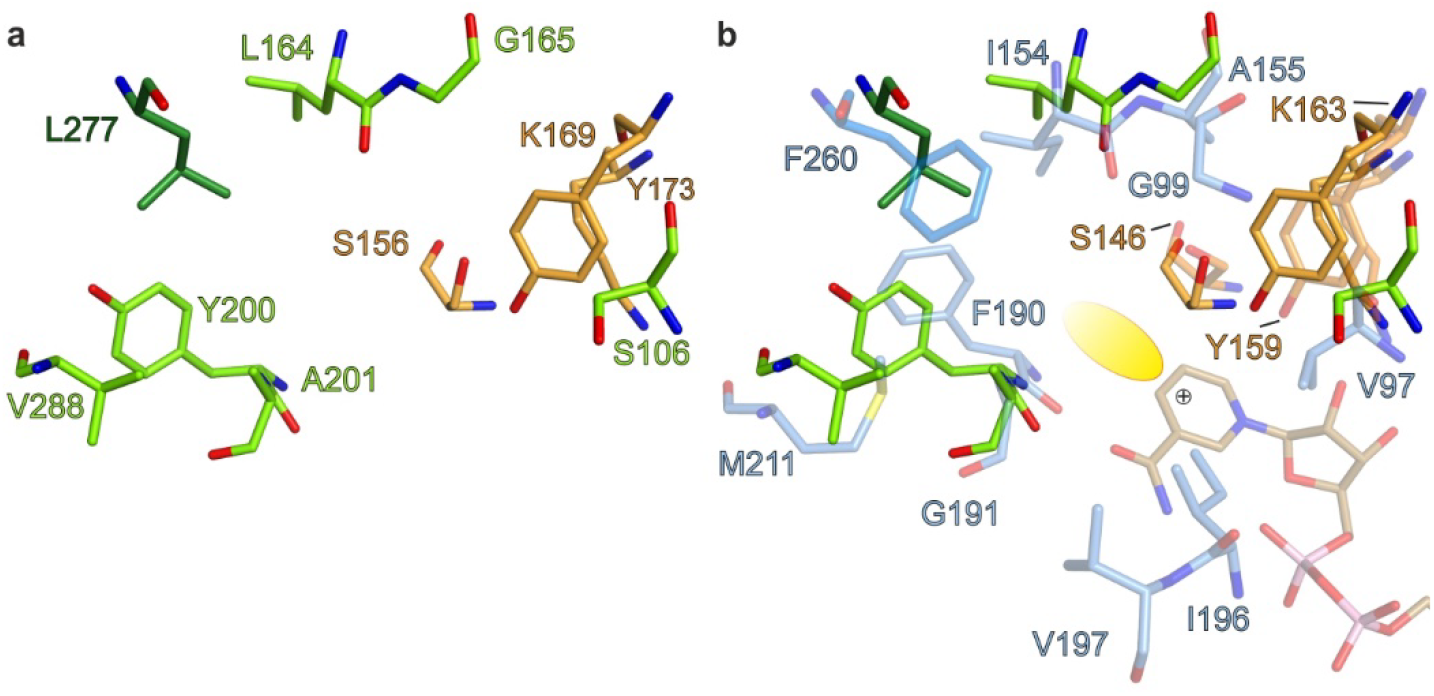
Active site architecture of *So*BDH2. Residues of the catalytic motif are colored in orange. **a** substrate binding pocket of *So*BDH2. Numbering refers to residues of *So*BDH2. **b** Superposition of *So*BDH2 and *Sr*BDH1•NAD^+^ (PDB ID 6ZYZ)^11^. Numbering of amino acids refers to *Sr*BDH1. Residues on equivalent position of I196 and V197 are not resolved in the density due to the absence of NAD^+^. Corresponding residues to latter two residues of *So*BDH2 are L206 and A207. The yellow ellipse indicates the potential substrate-binding site.

Due to fold differences, the active site architectures plant BDHs and *Ps*BDH differ drastically (**Supplementary Fig. 5c**). In both *So*BDH2 and *Sr*BDH1, the single αFG helix flanks the substrate-binding site, while the equivalent region in *Ps*BDH is divided into two discrete helices (**Supplementary Fig. 5c**), αFG1 and αFG2. Furthermore, the C-terminus of *Ps*BDH does not contribute to the substrate-binding site. The differences could be related to the natural functions of the enzymes. The plant BDHs are involved in the synthesis of the pure enantiomers of borneol and camphor as ingredients of essential oils, which requires high stereoselectivity. The bacterial enzyme, in contrast, participates in the catabolism of monoterpenols, where stereoselectivity does not provide an evolutionary advantage.

### Cryo-EM structure of *Sr*BDH1

To explore the general applicability of cryo-EM for the high-resolution structure analysis of small plant enzymes, we also subjected *Sr*BDH1 to cryo-EM-based structure analysis. The *Sr*BDH1 preparation yielded a single peak in a SEC/MALS analysis, consistent with a tetramer in solution, in agreement with its crystal structure^11^. As *Sr*BDH1 readily crystallized under various conditions, unlike *So*BDH2, we compared the thermal stability of both proteins by differential scanning fluorometry. Interestingly, the readily crystallizable *Sr*BDH1 is stabilized by approximately 8 °C compared to *So*BDH2 (**Fig. 2d**).

Cryo-grid preparation for *Sr*BDH1 was done as for *So*BDH2. To make sure that the resolution will not be limited by sampling of the detector, we decided to increase the magnification during data acquisition. By picking 1,635,690 particle images from 1,666 micrographs, we generated a dataset of similar size as for *So*BDH2. Following the same data processing routine as for *So*BDH2 yielded a final *Sr*BDH1 reconstruction at 1.88 Å resolution. (**Table 1, Supplementary Fig. 3** and **4**). Remarkably, the resolution of the cryo-EM structure of apo *Sr*BDH1 is much higher compared to the best resolved crystal structure *Sr*BDH1•NAD^+^ (PDB ID 6ZYZ)^11^, with four bound NAD^+^ molecules at 2.27 Å resolution. As assumed, *Sr*BDH1 is arranged as a tetramer (**Fig. 4** and **Supplementary Fig. 1**). During atomic modeling we followed the same refinement procedure as described for *So*BDH2 with the exception that we used the crystal structure of apo *Sr*BDH1 (PDB ID 6ZZ0)^11^ as starting model. The crystal structure and cryo-EM structure are practically identical (**Supplementary Table 3**). The density is of outstanding quality, allowing unambiguous assignment of amino acid side chains in alternate conformations (**Fig. 4**) and placement of water molecules.

**Fig. 4.**
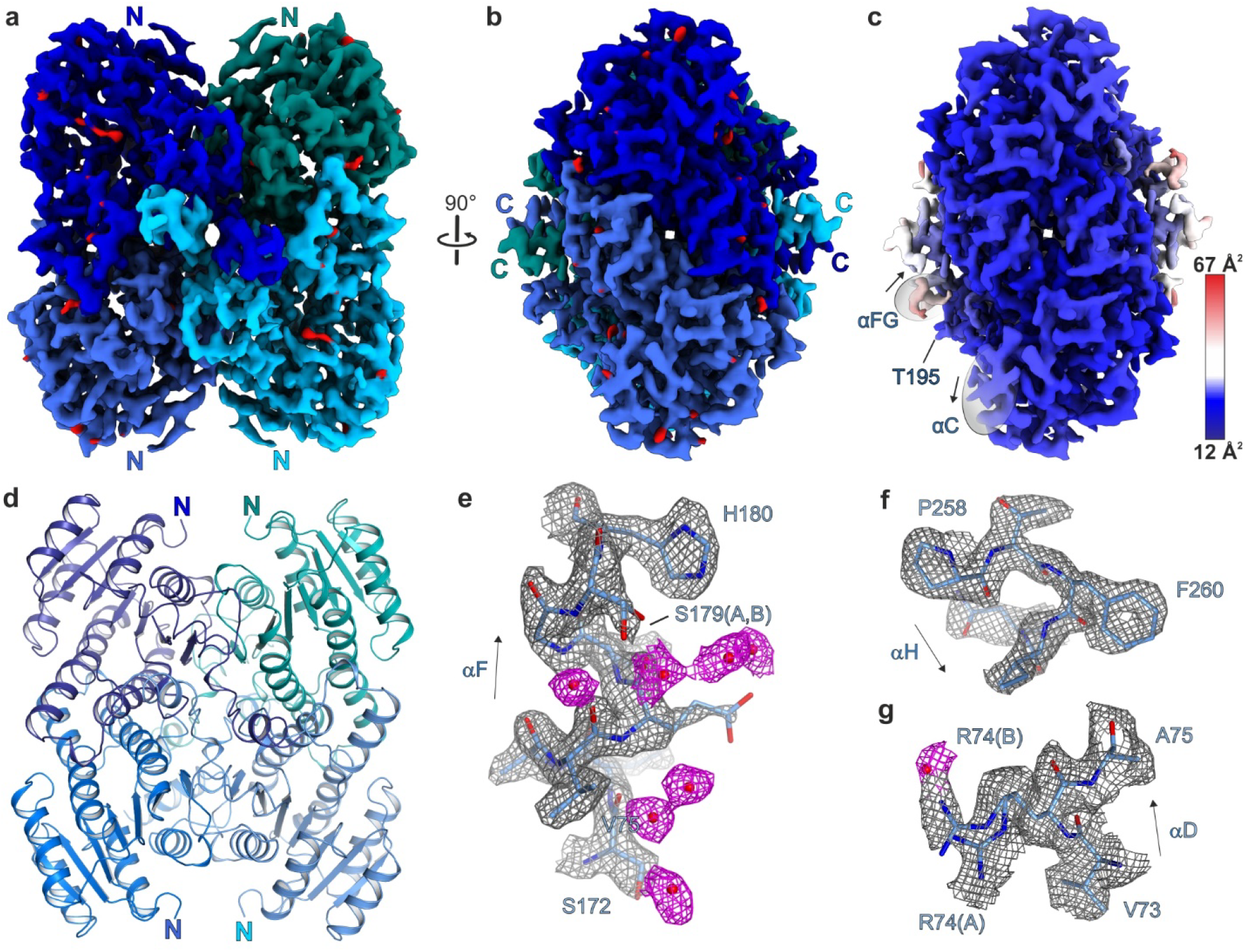
cryo-EM structure of *Sr*BDH1. **a** Tetrameric assembly of *Sr*BDH1. Density is shown for each of the four protomers in different blue shades. In red, locations of very well defined water molecules. **b** Same color-coding as in a, but rotated by 90 °. **c** Grouped B-factor mapped on the density. Color-gradient from blue to red, corresponding to increasing B-factors. In contrast to *So*BDH2, the B-factor distribution is uniform. **d** *Sr*BDH1 structure in cartoon representation. Same view and color-coding as in panel a. **e** Zoom on to the C-terminal end of αF with an alternate side chain conformation of S179 and well-defined water molecules, shown as red spheres. **f** C-terminal end of αH with F260 with a characteristic hole in the density for the aromatic ring system. **g** Double conformation of R74. The water molecule in distance of 2.5 Å to the guanidinium function of R74 in side chain conformation (B).

Almost the entire protein chain could be traced in the cryo-EM map. Apart from the first eight residues and the very C-terminal residue (**Fig. 2e**), the region from L193 to L205 is not defined in the density due to the missing NAD^+^ cofactor, as in *So*BDH2 (**Fig. 2e** and **Supplementary Fig. 5a**). In comparison to the available crystal structures of *Sr*BDH1, the total number of built residues is practically identical.

The *Sr*BDH1 model derived from the cryo-EM map is virtually identical to the apo-state crystal structure (r.m.s.d. of 0.6 Å for 982 pairs of Cα-atoms; **Supplementary Fig. 5b**). At 1.88 Å resolution, we could identify 399 water molecules, which uniformly cover the protein surface or are bound in cavities within the protein core. The ratio of water molecules to residues (0.4) is much lower compared to structures determined by X-ray crystallography with an expected number of one water molecule per residue at a resolution of 2.0 Å^26^. This discrepancy is explained by the absence of solvent channels in cryo-EM structures and missing local proximity of protein molecules. We observed 34 side chains with a double conformation, corresponding to about 3.5% of all residues. The observed ratio is perfectly in line with a detailed study describing that 3% of the residues present alternate side chain conformations in protein crystal structures with a resolution between 1.0 and 2.0 Å^27^.

Since the number of high-resolution cryo-EM structures is limited, we wondered whether the Ramachandran *Z*-scores^28^ of our structures (**Table 1**) would follow the distribution of Ramachandran *Z*-ranges as observed for crystal structures in a similar resolution regime^29^. The Ramachandran *Z*-scores of the *Sr*BDH1 and *So*BDH2 structures are in the expected region of crystal structures of similar resolution. Noteworthy, we refined the models without Ramachandran restraints, demonstrating that the Ramachandran Z-score can be a valuable measure for cryo-EM densities as well.

## Discussion

Structures of homomultimeric plant enzymes are under-represented in the fast-growing collection of protein structures analyzed by cryo-EM. Here we elucidated the cryo-EM structures of two comparatively small plant BDHs to high resolution. Given the molecular weight of the tetrameric complex, we report here on the highest resolution achieved by cryo-EM so far (**Supplementary Table 1**), pushing the boundaries of this rapidly developing method.

The new *So*BDH2 structure we describe revealed details of the enzyme’s active site architecture and allowed comparison to *Sr*BDH1. To our surprise, we could not observe NAD^+^ in the cryo-EM structure of *Sr*BDH1, although the protein sample used for crystallization and cryo-EM was identical. A possible explanation for this difference could be that in the crystal, the cofactor-binding loop is stabilized by crystal contacts and thus may have trapped NAD^+^. Alternatively, vitrification of the sample for cryo-EM may have led to the loss of the cofactor. We attempted to find an explanation for why *Sr*BDH1, but not *So*BDH2, could be crystallized. First, *Sr*BDH1 has a considerable higher Tm compared to *So*BDH2, suggesting a higher fold stability that may be more amenable to crystallization. Furthermore, although both cryo-EM structures superimpose with an r.m.s.d. of 1.3 Å for 952 pairs of Cα atoms (**Supplementary Fig. 5a**), local structural differences might have hindered crystallization of *So*BDH2. The αC helix of *So*BDH2 (Q52 to G65) is weakly defined in the density and hence much more flexible compared to *Sr*BDH1 (**Fig. 2c,e** and **3c**). Furthermore, in the *Sr*BDH1 structure, the αFG helix, upstream of the unresolved loop region, is stabilized by the C-terminal αH helix via hydrophobic contacts. In contrast, the C-terminus is shorter and not folded in an α-helix in *So*BDH2 (**Fig. 2c,e** and **Supplementary Fig. 5a**). Lastly, we cannot rule out that the NAD^+^ cofactor might stabilize *Sr*BDH1 to a larger extent, and its presence supports the crystallization process, which is not the case for *So*BDH2.

Given the small size of our protein samples and high particle density on the grids, sufficient data for high-resolution structure analysis could be rapidly acquired, reducing valuable instrument time. Given the high resolution of our structures, model building was greatly facilitated by automated routines, in particular ARP/wARP – ARPEM^20^ in combination with iterative refinement cycles in REFMAC5^30^. Moreover, due to the small protein size, real-space refinement and validation was fast.

During the past two decades, X-ray crystallography has been the main structural biochemical method to support drug development. Our observation that high-resolution (≤2.0 Å) structures of rather small proteins can be elucidated by cryo-EM in a short time emphasizes the important role that cryo-EM has to play in future drug development efforts, e.g., by using high-throughput applications such as fragment-based screening. Apart from circumventing time-consuming crystallization screening and possible phasing problems, an additional considerable advantage of cryo-EM in these and other endeavors is a much reduced sample consumption compared to crystallography. Likewise, our findings show that cryo-EM already today is an attractive tool for the structure analysis of enzymes used in green industry.

Availability of high-resolution structural data of newly discovered enzymes is crucial for understanding the molecular basis of their catalytic properties. Furthermore, with this knowledge, characteristics such as stability and selectivity can be improved by rational protein engineering instead of the time-consuming random mutagenesis approaches^31^. Rational design will greatly facilitate the generation of tailor-made enzyme in relatively short time periods. Cryo-EM is a valuable tool to achieve these goals, as it allows a fast and high-resolution structure determination for enzymes that proved difficult to crystallize.

## Methods

### Cloning

The synthetic genes of *So*BDH2 and *Sr*BDH1 borneol-type dehydrogenase from *S. officinalis* (GeneBank ID MT525099) and from *S. rosmarinus* (GeneBank ID MT857224) were ordered at GeneScript (USA), codon-optimized for *E. coli* and cloned into the vector pET15b in frame with an N-terminal His6-tag^11, 15^.

### *So*BDH2 expression and purification

*E. coli* BL21-RIL cells were transformed with a pET15a vector containing *So*BDH2 fused to an N-terminal hexa-histidine-tag. Protein induction was carried in auto-induction media at 37 °C for 7 h and subsequently cooled down to 16 °C^32^. Cells were grown over night and harvested by centrifugation (10 min, 7,000 rpm at 4 °C). The pellets were resuspended with 20 mM Tris/HCl pH 8.0, 500 mM NaCl (buffer A). Cells were lysed by homogenization at 4 °C for 7 min after addition of 0.5 mg l^−1^ DNase and the lysate was cleared by centrifugation (30 min, 21'500 rpm at 4 °C). A Ni^2+^-NTA column (cv 1 ml, Macherey Nagel) was equilibrated with buffer A and *So*BDH2 was loaded on the column and washed with 15 cv of buffer A. *So*BDH2 was eluted with buffer A supplemented with 300 mM imidazole. The protein was incubated with 3-fold molar excess of NAD^+^ (0.5 M in ddH2O) for 10 min on ice prior size-exclusion chromatography (SEC). SEC was performed with a HighLoad Superdex S200 16/60 column (GE Healthcare), equilibrated with 20 mM Tris/HCl, pH 8.0, 125 mM NaCl. Pooled protein fractions were concentrated with Amicon-Ultra-15 (Merck KGaA) to 11.2 mg ml^−1^ as measured by the absorbance at 280 nm. *Sr*BDH1 was purified by a practically identical protocol^11^.

### Size exclusion chromatography - multi-angle light scattering (SEC-MALS)

SEC-MALS experiments were performed at 18 °C. *So*BDH2 was loaded onto a Superdex 200 increase 10/300 column (GE Healthcare) that was coupled to a miniDAWN TREOS three-angle light scattering detector (Wyatt Technology) in combination with a RefractoMax520 refractive index detector. For calculation of the molecular mass, protein concentrations were determined from the differential refractive index with a specific refractive index increment (*d*n/*d*c) of 0.185 ml^−1^. Data was analyzed with the ASTRA 6.1.4.25 software (Wyatt Technology).

### Differential scanning fluorometry (DSF)

Melting temperature of proteins was measured with the Mx3005P qPCR system (Agilent) in a 96-well plate format under the buffer conditions used for crystallization or cryo-EM, respectively. Each well contained 10 μl buffer, 10 μl protein (0.15 μg μl^−1^) with 10x SYPRO Orange dye (Invitrogen) end concentration. The program consisted of three steps: step 1 was a pre-incubation for 1 min at 20°C, and steps 2 and 3 were cycles comprising the temperature increase of 1°C within 20 s. The temperature gradient proceeded from 25 to 95°C at 1°C per minute. Samples were measured in triplicates. The data was acquired with MxPro QPCR software (Agilent) and analysed with DSF Analysis v3.0.1 tool (ftp://ftp.sgc.ox.ac.uk/pub/biophysics) and Graphpad Prism 5.0.0.228 (Graph Pad Software Inc.). A t-test was performed with Graphpad Prism to validate the significance of the results.

### Cryo-electron microscopy

Samples were diluted to 1 mg/ml and a total of 3.8 μl was applied to glow-discharged 300 mesh holey gold UltrAuFoil R1.2/1.3 grids (Quantifoil Micro Tools GmbH). Vitrification was conducted using a Vitrobot Mark IV (Thermo Fisher Scientific, Eindhoven, Netherlands) set to 10 °C and 100% humidity by plunging into liquid ethane after 4 s blotting.

Data for the *So*BDH2 was collected on a FEI Titan Krios G3i transmission electron microscope (Thermo Fisher Scientific, Eindhoven, Netherlands) operated at 300 kV equipped with a Falcon 3EC at a nominal magnification of 96,000x, corresponding to a calibrated pixel size of 0.832 Å. Objective astigmatism and coma were corrected with AutoCTF (Thermo Fisher Scientific, Eindhoven, Netherlands) at final imaging conditions. To maximize beam coherence, a 50 μm C2 aperture was chosen. Direct alignments were executed thoroughly, beam parallelism and condenser astigmatism were optimized using the ronchigram on a volta phase plate (VPP), which was retracted during data acquisition. During imaging an electron flux of 0.7 e-/px*s on the detector was selected, corresponding to a dose rate of 1 e^−^/Å^2^* s on the sample. Images were taken at a nominal defocus between −0.6 and −1.6 μm accumulating a total dose of 40 e^−^/Å^2^ during a 40 s exposure, fractionated into 33 images. For automated data acquisition, EPU 2.8.1 (Thermo Fisher Scientific, Eindhoven, Netherlands) was utilized with AFIS enabled allowing for 6 μm image-beam-shift acquisition. The implemented ice filter was adjusted to exclusively image regions with thinnest ice.

Data for *Sr*BDH1 was acquired on the same instrument with minor exceptions. Nominal magnification was increased to 120,000x yielding a pixel size of 0.657 Å. The electron flux was adjusted to 0.6 e^−^/px*s on the detector resulting in a dose rate of 1.3 e^−^/Å^2*^s on the sample. During an exposure time of 31 s a total dose of 40 e^−^/Å^2^ was applied on the sample.

### Cryo-EM image processing

Raw movies of the *So*BDH2 dataset were aligned and dose-weighted with patch-motion correction implemented in cryoSPARC (2.9)^19^. Initial CTF estimation was achieved using Patch CTF. For initial particle picking the Blob picker was used with a particle diameter of 120-160 Å. Shiny class averages generated by reference-free 2D classification were selected as templates for template-based particle picking using a 120 Å circular mask. A total of 1,551,724 particle images were extracted with a box size of 224 px, fourier-cropped to 56 px (3.328 Å/px) for initial analysis and subjected to 40 iterations of 2D classification. Shiny classes were selected for *ab initio* reconstruction imposing *D*2 symmetry. Heterogeneous refinement with 3 classes didn’t guide further classification, therefore particle images were re-extracted fourier-cropped to a box size of 112 px (1.664 Å/px). The best resolved structure after heterogeneous refinement was re-extracted with a box size of 256 px (0.832 Å/px). NU refinement into a single class of 290,356 particles yielded a reconstruction with 2.32 Å resolution. Global and local CTF correction did not improve the resolution, however the reconstruction visually appeared better defined. In order to better account for anisotropic motion of the particles, local motion correction was applied, followed by Global CTF refinement yielding a reconstruction after NU refinement at 2.2 Å resolution. Micrographs with estimated resolutions worse than 3.5 Å were discarded, leaving 254,403 particle images for another cycle of local motion correction followed by Global CTF refinement and NU refinement. To account for point-spread of the signal in particle images, a box size of 384 px (320 Å) was used for re-extraction, giving a resolution after NU refinement of 2.1 Å. Another heterogeneous refinement run was conducted to isolate the final population of 173,781 particle images, which was reconstructed after local motion correction by NU-refinement to 2.0 Å resolution.

*Sr*BDH2 was refined similarly, with the exception that choosing the same final box size of 384 px resulted in smaller absolute dimensions of the box. From a total of 1587 micrographs 1,635,690 particle images were extracted, resulting in 410,573 selected particle images after reference-free 2D classification. After iterative homogeneous and heterogeneous refinement cycles, a final subset of 210,505 particle images was selected yielding a reconstruction with 1.88 Å resolution after NU refinement.

### Model building and refinement

An initial model of *So*BDH2 was obtained by automatic model building with ARP/wARP – ARPEM (8.0)^20^ using a sharpened electron density volume and the protein sequence as input. Automatic model building comprised iterative refinement in REFMAC5 (5.8.0258)^30^. For comparison, the phenix.map_to_model procedure^21^ as well as BUCCANEER^22^ as part of the CCPEM suite^33^ were used for automated model building. The obtained model was manually adjusted to the electron density, supported by real space refinement in COOT (0.8.9.1)^23^. The model was refined against the cryo-EM map using the real space refinement protocol in PHENIX (1.19.1)^34, 35^. Water molecules were added in COOT and manually inspected, followed by an additional round of real space refinement in PHENIX. In final stages of refinement, we fully released the restraints for secondary structure elements, Ramachandran, non-crystallographic symmetry (NCS), and no corrections of energetically disfavored rotamer conformations. In final rounds of refinement, grouped atomic displacement factors were refined. The structures were evaluated with EMRinger^36^ and MOLPROBITY^37^. Structure figures were prepared using PyMOL (Version 1.8 Schrödinger, LLC) and CHIMERA^38^. Secondary structure elements were assigned with DSSP^39^ and ALSCRIPT^40^ was used for secondary structure-based sequence alignments.

## Data availability

The atomic models have been deposited in the Protein Data Bank (PDB) with the following accession codes: 2.04 Å structure of *So*BDH2 with PDB ID 7O6P, and 1.88 Å structure of *Sr*BDH1 with PDB ID 7O6Q. The cryo-EM maps have been deposited in the Electron Microscopy Data Bank as follows: 2.04 Å map (EMD-12739), 1.88 Å map (EMD-12740). All other data supporting the findings of this study are available from the corresponding author on request.

## Acknowledgement

The authors acknowledge financial support by the German Federal Ministry of Education and Research (BMBF) for the project CbP-camphor based polymers within the bio-economy international program (grant No. 031B050B). R.K. and A.C. also would like to thank the Austrian Science Funds (FWF, P31001-B29) for financial support. We acknowledge access to electron microscopic equipment at the core facility BioSupraMol of Freie Universität Berlin, supported through grants from the Deutsche Forschungsgemeinschaft (HA 2549/15-2), and from the Deutsche Forschungsgemeinschaft and the state of Berlin for large equipment according to Art. 91b GG (INST 335/588-1 FUGG, INST 335/589-1 FUGG, INST 335/ 590-1 FUGG). C.P.O. Helmer is supported by the Hanns Seidel Foundation. We acknowledge technical support in protein purification by C. Langner and support by Y. Huang with the SEC/MALS experiments. The authors like to thank B. Kirmayer and B. Schade for assistance with cryo-EM sample preparation and microscope operation.

## Contributions

A.M.C. cloned the construct and made expression tests. C.P.O.H. produced protein samples with help from N.D. and B.L. N.D. perfomed SEC/MALS analyses and prepared cryo-EM samples with T.H., built atomic models with B.L. and M.C.W. and refined structures. T.H., acquired, processed and refined cryo-EM data. All authors contributed to the analysis of the data and the interpretation of the results. B.L., T.H., and R.K. wrote the manuscript with contributions from the other authors. B.L. and R.K. supervised work in their respective groups. B.L., T.H., and R.K. conceived and coordinated the project.

