## Supplementary Information for "Rapid high-resolution structure analysis of small, biotechnologically relevant enzymes by cryo-electron microscopy"

Dimos *et al.*

**Table of content:**

**Supplementary Table 1.** Overview of selected cryo-EM structures with the highest achieved resolution.

**Supplementary Table 2.** Comparison of the performance of different automated model building programs.

**Supplementary Table 3.** Structural comparison of the three different crystal structures of *Sr*BDH1.

**Supplementary Fig. 1**. Cryo-EM analysis of *So*BDH2.

**Supplementary Fig. 2.** Data processing workflow for the *So*BDH2 dataset.

**Supplementary Fig. 3**. Cryo-EM analysis of *Sr*BDH1.

**Supplementary Fig. 4.** Data processing workflow for the *Sr*BDH1 dataset.

**Supplementary Fig. 5.** Structural comparison of related BDH structures focusing on the substrate/cofactor binding site.

**References**

**Supplementary Table 1.** **Overview of selected cryo-EM structures with the highest achieved resolution.** The summary does not include larger multi-subunit complexes. Membrane proteins incorporated in nanodiscs are indicated with an asterix.

| **enzyme** | **organism** | **assembly** | **resolution**  **[Å]** | **total M_r_ [kDa]** | **EMDB**  **ID** | **reference** |
| --- | --- | --- | --- | --- | --- | --- |
| apoferritin | *H. sapiens* | 24-mer | 1.15 | 480 | 11668 | ^1^ |
| β_3_ GABAA receptor* | *H. sapiens* | pentamer | 1.7 | 200 | 11657 | ^2^ |
| β-galactosidase | *E. coli* | tetramer | 1.8 | 465 | 21995 | ^3^ |
| BDH1 | *S. rosmarinus* | tetramer | 1.88 | 120 | 12740 | This study |
| urease | *H. pylori* | dodecamer | 2.04 | 1100 | 11233 | ^4^ |
| BDH2 | *S. officinalis* | tetramer | 2.04 | 129 | 12739 | This study |
| ORF3a/apolipo* | SARS-CoV-2 | tetramer | 2.08 | 114 | 22898 | ^5^ |
| CDK-activating kinase | *H. sapiens* | dimer | 2.51 | 119 | 12042 | ^6^ |
| aldolase | *O. cuniculus* | tetramer | 2.6 | 150 | 8743 | ^7^ |
| catalase-peroxidase | *M. tuberculosis* | dimer | 2.68 | 161 | 11776 | ^8^ |
| alcohol dehydrogenase | *S. carlsbergensis* | tetramer | 2.7 | 147 | 22807 | ^9^ |
| methemoglobin | *H. sapiens* | tetramer | 2.8 | 64 | 0407 | ^10^ |
| lactate dehydrogenase | *G. gallus* | tetramer | 2.8 | 144 | 8191 | ^11^ |
| alcohol dehydrogenase | *E. caballus* | dimer | 2.9 | 82 | 0406 | ^10^ |
| biotin-bound streptavidin | *S. avidinii* | tetramer | 3.2 | 52 | 0689 | ^12^ |
| cytotoxin A | *H. pylori* | hexameric | 3.2 | 530 | 0542 | ^13^ |
| isocitrate dehydrogenase | *H. sapiens* | dimer | 3.8 | 93 | 8193 | ^11^ |
| catalytic subunit protein kinase A | *M. musculus* | monomer | 6.0 | 43 | 0409 | ^10^ |

**Supplementary Table. 2.** **Comparison of the performance of different automated model building programs**. ARP/wARP – ARPEM^14^, phenix.map_to_model^15^, as well as Buccaneer^16^. Green numbers refer to the structure of *So*BDH2 and blue numbers to *Sr*BDH1.

|  | **ARP/wARP**  **ARPEM** | **Phenix**  **map_to_model** | **CCPEM**  **Buccaneer** | **Final model** |
| --- | --- | --- | --- | --- |
| Total number of residues | **1212** / **1160** | | | |
| Residues built | **1001 / 963** | **880 / 828** | **1159 / 972** | **1022 / 977** |
| Residues sequenced | **922 / 921** | **880 /** | **1047 / 955** | **1022 / 977** |
| Completeness by residues built [%] | **82.0 / 83.0** | **72.6 /** | **91.5 / 83.8** | **84.5 / 84.2** |

**Supplementary Table 3.** **Structural comparison of the three different crystal structures of *Sr*BDH1.** *Sr*BDH1 apo (PDB ID 6ZZ0), *Sr*BDH1•NAD^+^ / high salt (PDB ID 6ZZT) with one bound NAD^+^ as well as *Sr*BDH1•NAD^+^ / PO/OH (PDB ID 6ZYZ) with four bound NAD^+^ molecules ^17^ as well as the two cryo-EM structures of *Sr*BDH1 and *So*BDH2. R.m.s.d. for pairs of Cα-atoms calculated with SSM^18^ as implemented in COOT^19^.

|  | ***Sr*BDH1 apo** | ***Sr*BDH1•NAD^+^ / high salt** | ***Sr*BDH1•NAD^+^ / PO/OH** | **cryo-EM *Sr*BDH1** | **cryo-EM *So*BDH2** |
| --- | --- | --- | --- | --- | --- |
| ***Sr*BDH1 apo** |  |  |  |  |  |
| ***Sr*BDH1•NAD^+^ / high salt** | 0.48 |  |  |  |  |
| ***Sr*BDH1•NAD^+^ / PO/OH** | 0.56 | 0.52 |  |  |  |
| **cryo-EM *Sr*BDH1** | 0.57 | 0.49 | 0.39 |  |  |
| **cryo-EM *So*BDH2** | 1.44 | 1.39 | 1.39 | 1.15 |  |

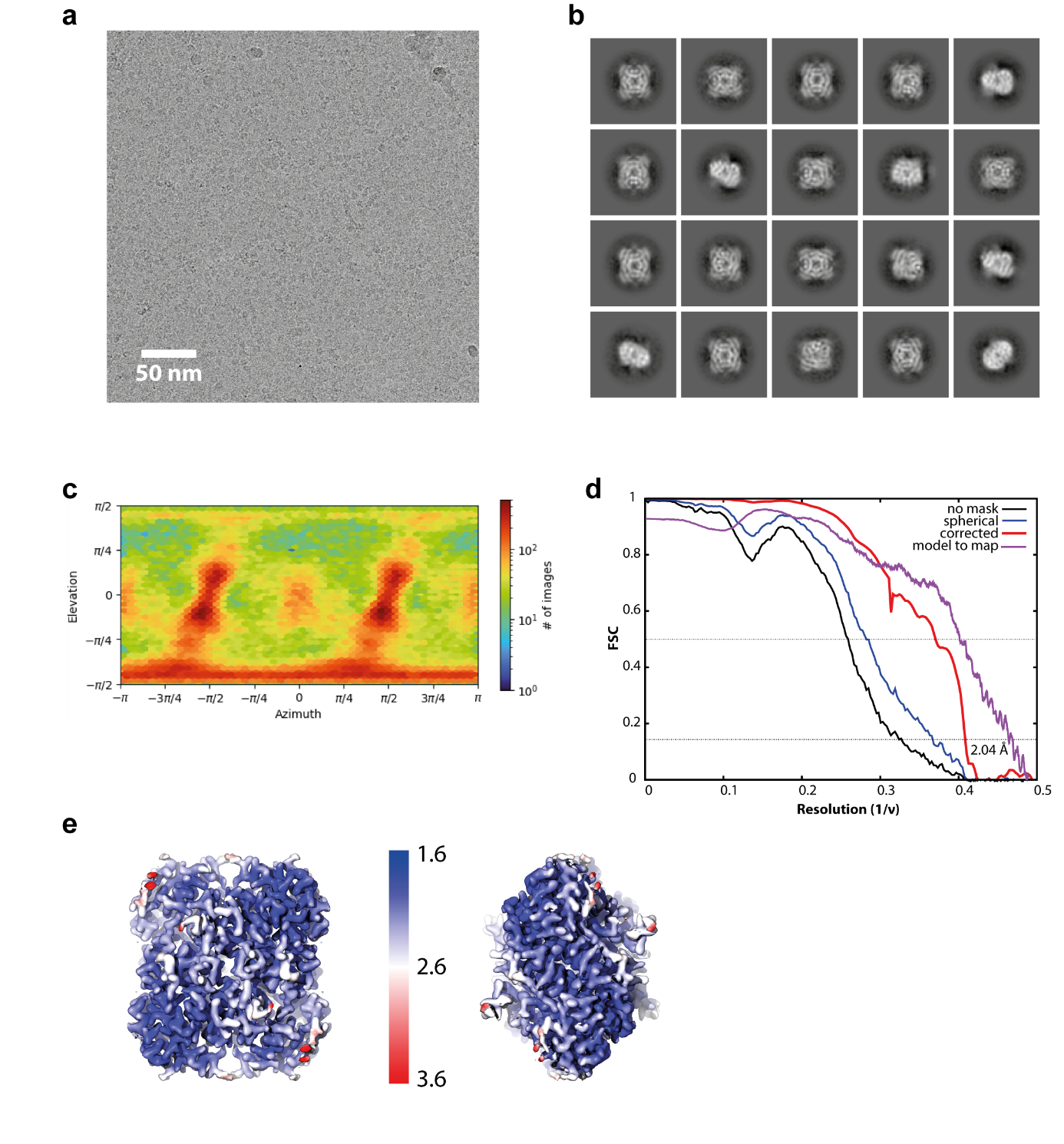

**Supplementary Fig. 1 Cryo-EM analysis of *So*BDH2. a** represents a representative cryo-EM micrograph, the scale bar indicates 50 nm spacing. **b** selected 2D class averages after reference-free 2D classification with cryoSPARC. Top and side views can be identified, excluding preferential orientation issues. A circular mask of 120 Å diameter was used during classification. **c** Viewing direction distribution as determined during non-uniform refinement with cryoSPARC. **d** Resolution estimates by fourier-shell correlation using either no mask (black line), a generous spherical mask (blue line) and after solvent correction by phase randomization (red line). Dashed lines represent FSC(0.5) and FSC(0.143) crossings. Model to map correlation as determined with PHENIX is colored purple. **e** Illustration of the local resolution estimation calculated with cryoSPARC for two different views of *So*BDH2 after rotation by 90°. Coloring of the cryo-EM density reflects the local resolution ranging from 1.6 Å to 3.6 Å. A major fraction of the structure is resolved well beyond 2 Å, less resolved regions are mainly situated in the periphery of *So*BDH2.

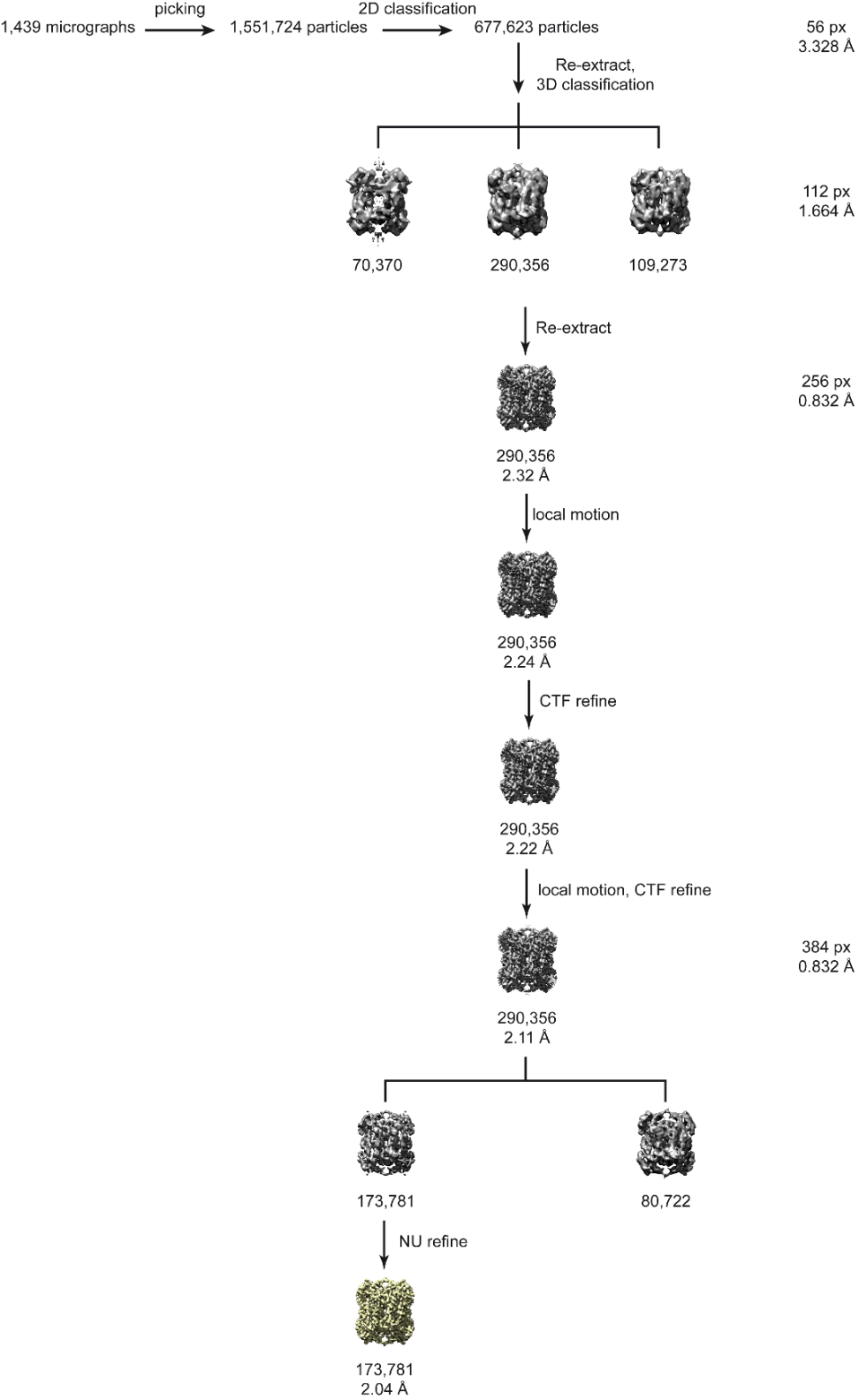

**Supplementary Fig. 2 Data processing workflow for the *So*BDH2 dataset.** From initially selected 1,439 micrographs ~1.5 M particles were picked and subjected to reference-free 2D classification. ~678k particle images were re-extracted with a box-size of 224 px fourier-cropped to 112 px giving a pixel-size of 1.664 Å. After 3D classification, a subset of 290,356 particle images was again re-extracted at full resolution with a larger box size of 256 px and homogeneously refined to 2.32 Å resolution. Particle based local motion correction improved the resolution to 2.24 Å, which could be only marginally improved to 2.22 Å by CTF refinement. Another cycle of local motion correction was applied using a larger extraction box of 384 px to preserve high frequency information of the CTF. Following CTF refinement the resolution improved to 2.11 Å. By heterogeneous refinement, a final subset of 173,781 particle images was selected for homogeneous NU refinement, yielding the final reconstruction at 2.04 Å resolution.

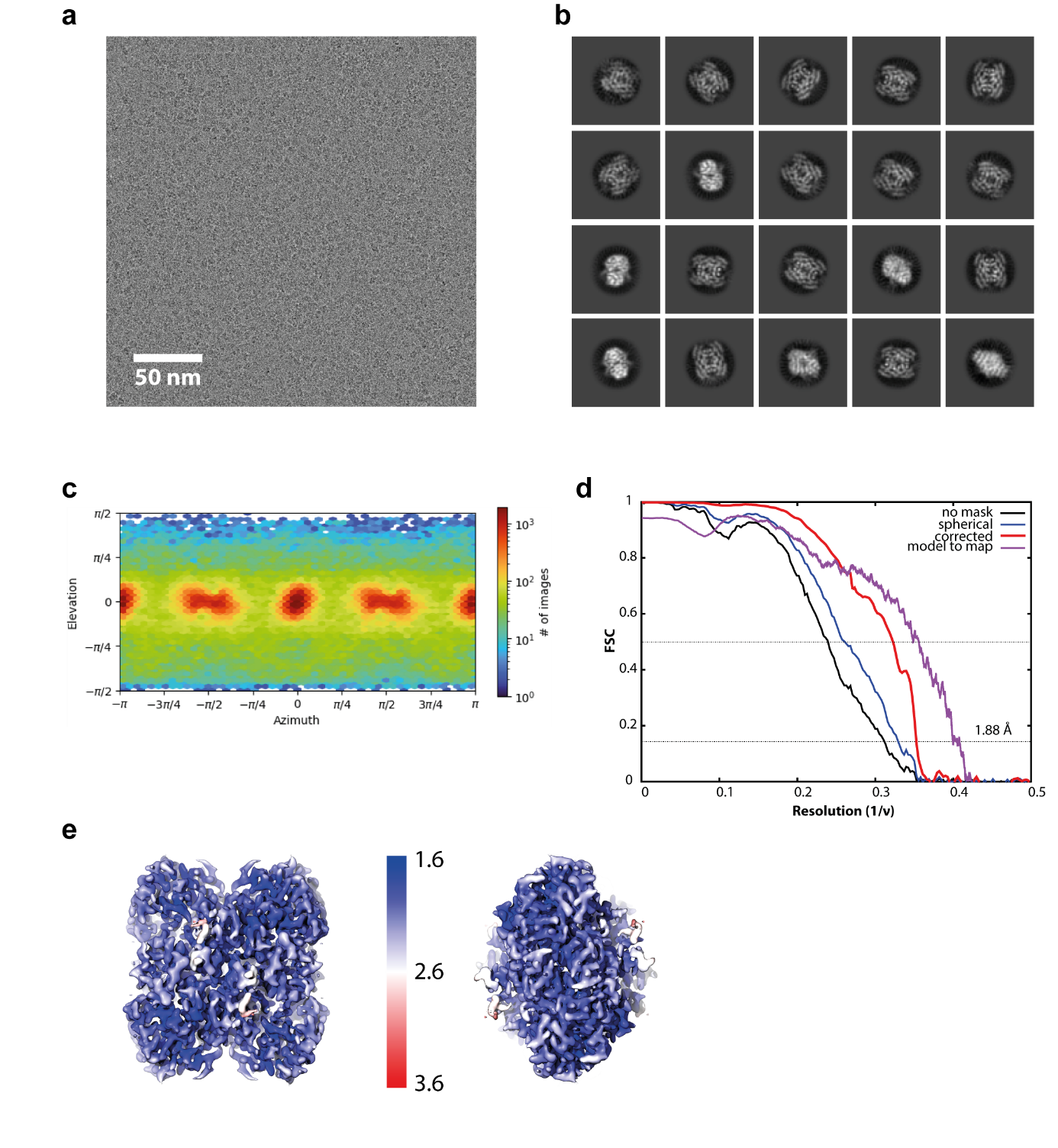

**Supplementary Fig. 3 Cryo-EM analysis of *Sr*BDH1. a** Representative cryo-EM micrograph, the scale bar indicates 50 nm spacing. **b** selected 2D class averages after reference-free 2D classification with cryoSPARC. As for *So*BDH2, top and side views can be identified. A circular mask of 100 Å diameter was used during classification. **c** Viewing direction distribution as determined during non-uniform refinement with cryoSPARC. **d** Resolution estimates by fourier-shell correlation using either no mask (black line), a generous spherical mask (blue line) and after solvent correction by phase randomization (red line). Dashed lines represent FSC(0.5) and FSC(0.143) crossings. Model to map correlation as determined with PHENIX is colored purple. **e** Illustration of the local resolution estimation calculated with cryoSPARC for two different views of *Sr*BDH1 after rotation by 90°. Coloring of the cryo-EM density reflects the local resolution ranging from 1.6 to 3.6 Å. The vast majority of the structure is resolved well beyond 2 Å.

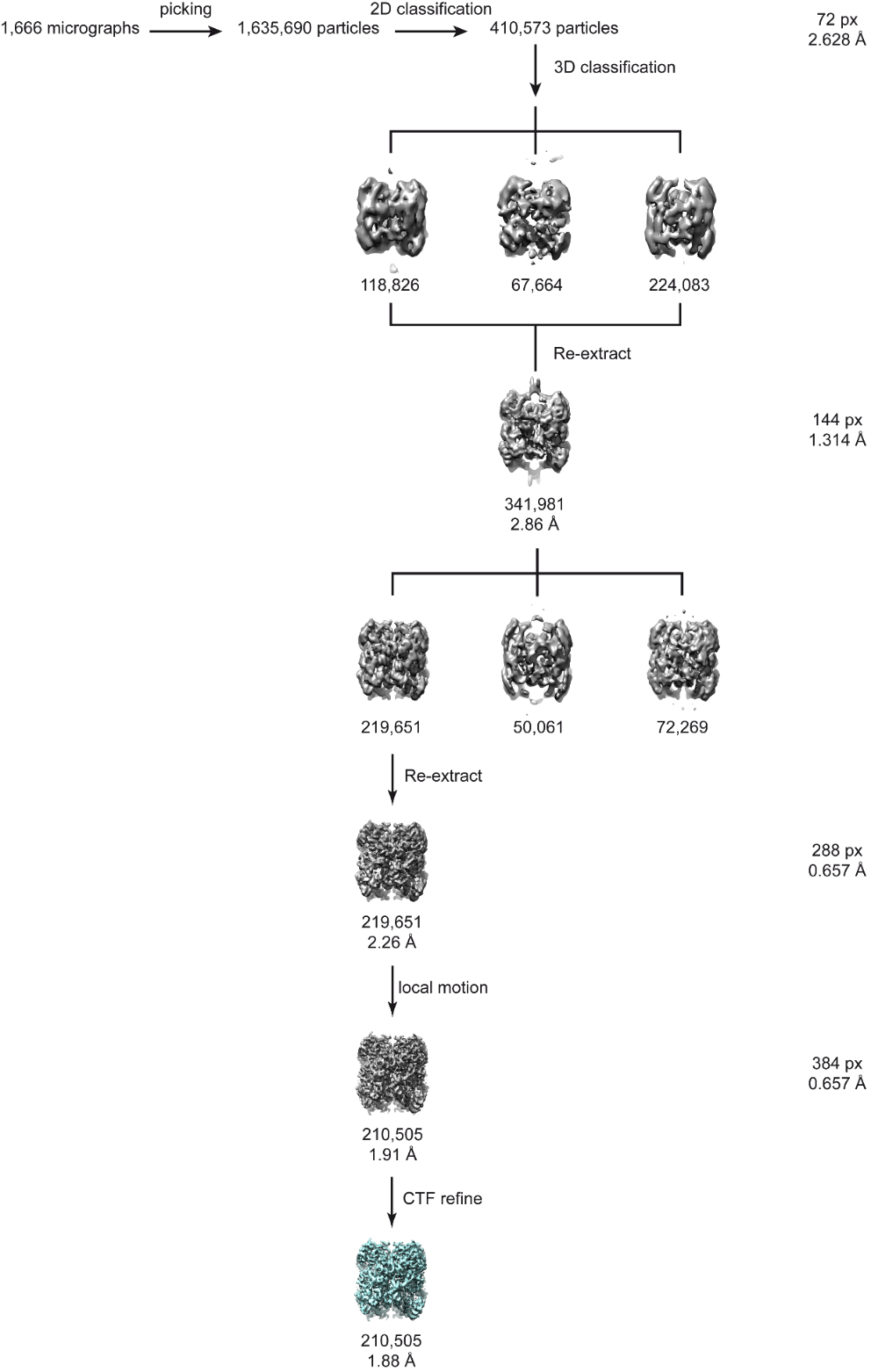

**Supplementary Fig. 4 Data analysis of the *Sr*BDH1 dataset.** Using the *So*BDH2 structure as reference, ~1.6 M particles were automatically picked from 1,666 micrographs with cryoSPARC. Iterations of 2D classification were applied to select 410,573 particle images for heterogeneous 3D classification. A subset of 341,981 was re-extracted with a box-size of 288 px, fourier-cropped to 144 px and homogeneously refined to 2.86 Å resolution. After another heterogeneous refinement using 3 classes, 219,651 particles were re-extracted at full resolution (0.657 Å/pix) yielding a reconstruction of 2.16 Å. Local motion correction was applied, after which 210,505 particles were re-extracted with a box size of 384 px and homogeneously refined to 1.91 Å resolution. CTF refinement followed by NU refinement generated the final reconstruction with 1.88 Å resolution (cyan).

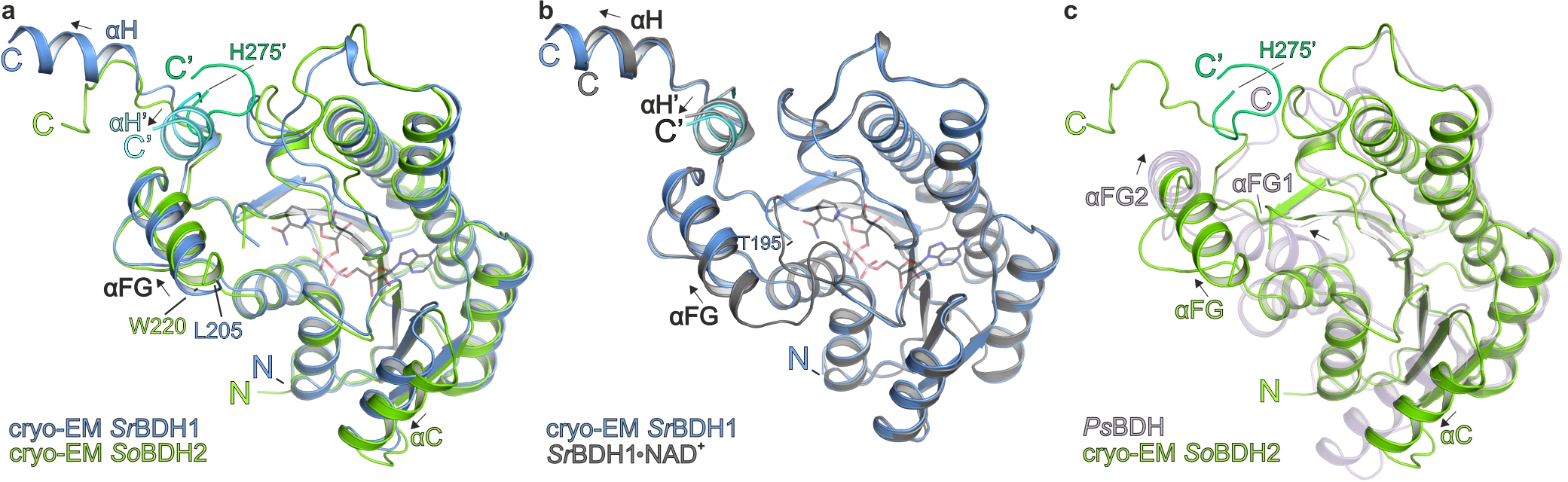

**Supplementary Fig. 5 Structural comparison of related BDH structures focusing on the substrate/cofactor binding site.** Only one protomer of the tetrameric complexes is shown. The proteins are shown in cartoon representation. **a** Superposition of the cryo-EM structures of *Sr*BDH1, drawn in blue, as well as *So*BDH2, drawn in green cartoon. The C-terminal αH helix of another protomer completes the substrate binding site. The NAD^+^ molecule is drawn in black, obtained by a superposition with the crystal structure of *Sr*BDH1•NAD^+^ / PO/OH (PDB ID 6ZYZ) ^17^. Structural differences can be seen in particular for the C-terminus and the αC helix. **b** Superposition of the cryo-EM structure of *Sr*BDH1 drawn in blue and the crystal structure of *Sr*BDH1•NAD^+^ / PO/OH (PDB ID 6ZYZ^17^) drawn in gray.Binding of NAD^+^ leads to stabilization of the loop region upstream of helix αFG and the helix itself. **c** Superposition of the cryo-EM structure of *So*BDH2 and the crystal structure of *Ps*BDH PDB ID 6M5N^20^) shown in light purple. Major structural differences are observed for the C-terminal portion of the protein.
